# Exploring Relationships between Functional Network Connectivity and Cognition with an Explainable Clustering Approach

**DOI:** 10.1101/2022.07.23.501266

**Authors:** Charles A. Ellis, Martina Lapera Sancho, Robyn Miller, Vince Calhoun

## Abstract

The application of clustering algorithms to fMRI functional network connectivity (FNC) data has been extensively studied over the past decade. When applied to FNC, these analyses assign samples to an optimal number of groups without a priori assumptions. Through these groupings, studies have provided insights into the dynamics of network connectivity through the identification of different brain states and have identified subgroups of individuals with unique brain activity. However, the manner in which underlying brain networks influence the identified groups is yet to be fully understood. In this study, we apply k-means clustering to resting-state fMRI-based static FNC data collected from 37,784 healthy individuals. We identified 2 groups of individuals with statistically significant differences in cognitive performance in several test metrics. Then, by applying two different versions of G2PC, a global permutation feature importance approach, and logistic regression with elastic net regularization, we were able to identify the relative importance of brain network pairs and their underlying features to the resulting groups. Through these approaches, together with the visualization of centroids’ connectivity matrices, we were able to explain the observed differences in cognition in terms of specific key brain networks. We expect that our results will shed further light upon the effect of underlying brain networks on encountered cognitive differences between groups with unique brain activity.

## I. Introduction

Clustering-based analyses have become increasingly common for the domain of resting-state functional magnetic resonance imaging (rs-fMRI) functional network connectivity (FNC) analysis. They have been used to identify patient subgroups or transitory states in brain activity associated with neurological or neuropsychiatric disorders or cognitive functions. While the identification of these subgroups and states have enabled a better understanding of various neurological [1] conditions and cognitive functions [2], existing analyses have not included explainability approaches for insight into what FNC values are actually most impactful upon the clustering and most likely to be associated with neurological or neuropsychiatric disorders and cognitive functions. In this study, we use several explainability approaches to gain insight into key brain networks associated with differences in cognitive function identified with a clustering approach.

FNC measures the interaction of brain regions and networks. FNC can be static (sFNC) indicating the average correlation across a whole resting state recording or dynamic (dFNC) demonstrating correlation in multiple time intervals [3]. A number of methods have been used for FNC analysis, including traditional statistical testing, supervised machine learning classification [4], and clustering [3].

Clustering of FNC data has become an increasingly common analysis approach. When sFNC is used during clustering, novel subgroups of individuals can be identified, and when dFNC is used, transitory states of brain activity associated with cognitive function [2] and neurological disorders [1][3] can be identified. Once subgroups or states are identified, cluster centroids are often compared to identify the interactions between brain networks that are most influential to the clustering. However, clustering explainability approaches that can identify the relative impact of each network pair upon the clustering are generally not applied.

In this study, we analyze a large whole-brain sFNC dataset from nearly 38,000 healthy individuals to identify subgroups of individuals with differences in cognitive performance. We then apply and compare several clustering explainability approaches for insight into the brain network pairs that most influence the identified clusters. One of the explainability approaches that we include represents a novel variation of Global Permutation Percent Change (G2PC) feature importance, a clustering explainability method, that provides better insight into the importance of each connectivity pair within networks.

## II. Methods

### A. Description of Dataset

Our dataset was composed of 37,784 samples of rs-fMRI and cognitive score data from the UK BioBank [5]. rs-fMRI data was collected with a Siemens Skyra 3T with a standard Siemens 32-channel RF receive head coil. A Gradient Echo EPI sequence with 8x multislice acceleration, no iPAT, a flip angle of 52^°^, and with fat saturation was used. Recordings were for 6 minutes (voxel size = 2.4×2.4×2.4 mm, TR = 0.735, TE = 39ms).

The cognitive function of the participants was assessed with 4 tasks [6]. (1) Numeric Memory task scores included the maximum number of correctly remembered digits (NM1) and time to complete test (NM2). (2) Fluid Intelligence task scores included the participant’s score (FI1) and the number of questions attempted within the time limit (FI2). (3) The Pair Matching task included the times to complete three distinct rounds with subsequent higher numbers of cards to be remembered (i.e., PM1, PM2, and PM3). Those individuals that did not complete a round did not proceed to the next round. (4) Prospective Memory task scores included the result for the test (PrM1) and time to answer (PrM2). Because PrM2 was measured differently across study sites, we excluded it from the analysis. The gender of the patients was also included.

### B. Description of Data Preprocessing

After fMRI data collection, we spatially normalized the data to the standard Montreal Neurological Institute (MNI) space with an EPI template with 3×3×3 mm^3^ isotropic voxels. We next smoothed the data with a Gaussian kernel with a full width at half maximum of 6 mm. We then used the Neuromark fully automated independent component analysis pipeline to extract 53 independent components (ICs) associated with 7 brain networks. The 7 networks included the subcortical (SCN), auditory (ADN), sensorimotor (SMN), visual (VSN), cognitive control (CCN), default mode (DMN), and cerebellar (CBN) networks. We then calculated the Pearson’s correlation between each of the ICs, resulting in 1,378 static functional network connectivity (sFNC) features that could be divided into 28 network pairs. The sFNC between two networks can be abbreviated as SCN/CBN in the case of the SCN and CBN networks.

Most study participants were missing one or more cognitive test scores. To avoid undue bias that might result from interpolating missing values, missing test scores were not included in our statistical analysis.

### C. Description of Clustering Approach

We used k-means clustering to partition the sFNC data into clusters. The optimal number of clusters (k=2) was determined through the Elbow and Silhouette methods using the Euclidean distance metric. Algorithm parameters (maximum number of iterations and number of initializations) were subsequently adjusted until convergence of results was reached (convergence was reached before default values; at 300 maximum iterations and 10 initializations).

### D. Description of Explainability Approaches

After assigning data for each of the subjects to different clusters, we applied two approaches to gain insight into the key brain network pairs differentiating the two clusters. We used G2PC Feature Importance, a recently developed explainability method that is broadly applicable across clustering algorithms, and we trained a logistic regression classifier with elastic net regularization (LR-ENR) with interpretable weights.

#### 1) G2PC Feature Importance

G2PC was first developed in [7], as an explainability approach that is easy to implement and that can provide explanations for many different clustering algorithms. In effect, it measures the sensitivity of clusters to the perturbation of different features or groups of features with the assumption that those features to which clusters are more sensitive are more important. G2PC has several steps: (1) data is assigned to clusters, (2) one feature or group of features is permuted across samples, (3) the percent of samples that switch clusters is calculated, and (4) the permutation is repeated for each feature or group of features m times. In our implementation, we used each of the 28 network pairs as a group during the permutation, as a previous study showed that perturbing one of the 1,378 sFNC features does not have a significant enough effect on cluster assignments to be picked up by G2PC. We permuted each group of features 100 times. Additionally, we realized that the original implementation of G2PC (G2PCu) did not account for the number of features in each group, meaning that larger groups would tend to have more of an effect upon the clustering even if some of the features in the group were of relatively little importance to the clustering. As such, we normalized the importance of each network pair by dividing by its corresponding number of features (G2PCn).

#### 2) Interpretable LR-ENR Classifier

To ensure that we identified network pairs truly influential to the clustering, we used a second explainability approach for comparison to G2PC. Specifically, we trained a LR-ENR classifier with 10-fold nested cross validation using a maximum number of iterations of 200,000 and several L1 ratios (0.25, 0.5, 0.75, 0.85, 0.9, 0.95, 0.99). Training, validation, and test groups composed 64%, 16%, and 20% of the data in each fold, respectively. LR-ENR is an interpretable model, such that the weight associated with each feature can be easily examined. To calculate the effect of each network pair upon the classification, we multiplied the magnitude of each model weight by the magnitudes of its corresponding feature values across all samples. We then averaged the resulting values across samples and averaged across the features in each network pair.

### E. Description of Statistcal Analyses for Cognitive Scores

After clustering, we sought to understand whether there were significant differences in the cognitive performance of the individuals in each group. As such, we applied several statistical analyses to the preprocessed cognitive performance data.

We performed Chi-squared tests for sex and PrM1, which were categorical, and normality tests for numerical scores followed by two-tailed Mann-Whitney U tests since normality test results showed that data was not normally distributed. When there were significant differences, we compared the U-statistics for each group to determine which group had higher performance for each measure. For all statistical tests, we used p < 0.05 to determine significance.

## III. Results and Discussion

In this section, we describe and discuss the results of our clustering and cognitive score analyses. We further discuss some of the limitations and next steps related to our work.

### A. Identification of Two Groups with Distinct FNC

Two well-defined clusters (referred to as groups 0 and 1) with a large difference in size were identified. Group 0 (G0) was much smaller than group 1 (G1) and had 9,967 individuals. In contrast, group 1 had 27,817 individuals. As shown in Figure 1A and 1B, to better understand what differentiated the two groups, we first examined the connectivity values associated with the centroids of each cluster.

**Figure 1.**
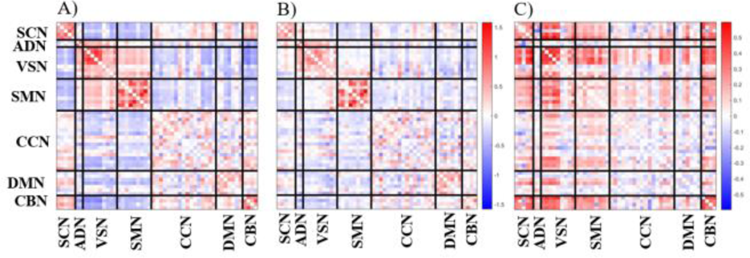
sFNC Matrices. A) Cluster centroid for G0. B) Cluster centroid for G1. C) Subtraction of absolute connectivity values of centroid 1 from absolute connectivity values of centroid 0. It should be noted that while the term ‘connectivity’ generally applies to positive and negative values, panel C solely focuses on the magnitude of these connections. The color bar to the right of panel B is for both panels A and B.

Interestingly, the magnitude of connectivity values in G0 were generally larger than those of G1. To further emphasize this difference, we subtracted the absolute values of centroid 1 from those of centroid 0 (Figure 1C). Interestingly, the connectivity values between the majority of feature pairs are above 0, indicating that absolute connectivity in G0 overall exceeds that of G1. Eleven brain networks had particularly prominent differences in connectivity between the clusters: CBN, CBN/SCN, SMN/SCN, CBN/SMN, SMN, VSN/SMN, ADN/SCN, SCN, CBN/VSN, CBN/ADN and SCN/VSN.

As shown in Figure 2, these results align with the results provided by G2PCn and LR-ENR Both methods, due to normalization of group feature importance (i.e., in G2PC) and averaging of importance across features (i.e., in LR-ENR), indicate the importance of each underlying feature within each network pair. As observed, both methods find that the 10 brain networks with most influential features are CBN, CBN/SCN, SMN/SCN, CBN/SMN, SMN, VSN/SMN, ADN/SCN, SCN, CBN/VSN, CBN/ADN. These coincide with the networks with the highest connectivity differences between groups. Additionally, both LR-ENR and G2PCn indicate that features within the CBN had the highest effect on the clustering.

**Figure. 2.**
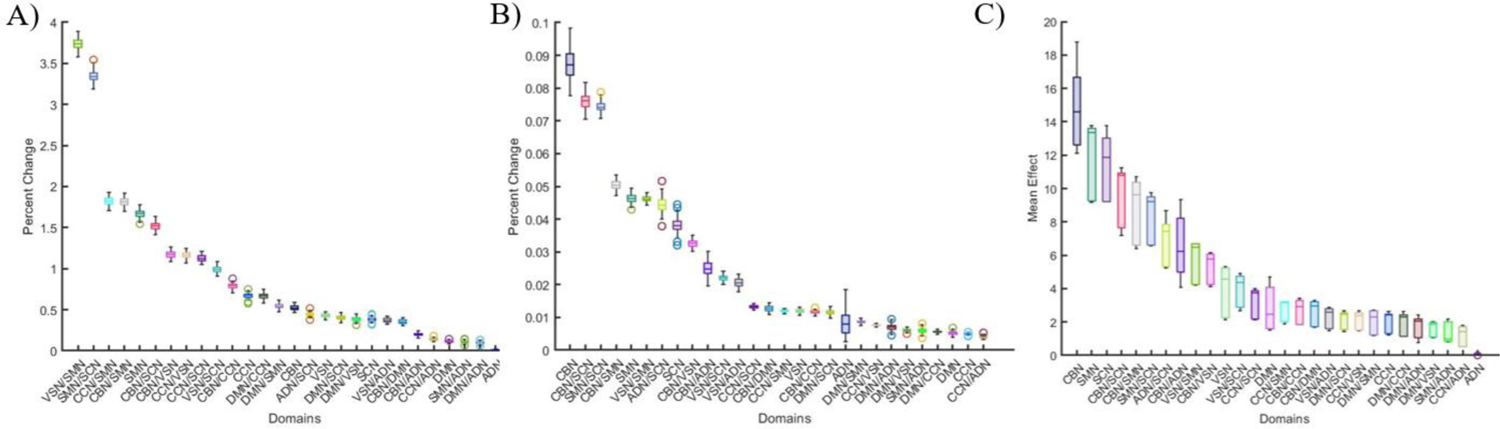
Importance of brain network pairs. A) G2PCu results. B) G2PCn results. C) LR-ENR results. The x-axes indicate each domain ordered based upon their median importance from left to right. The y-axes indicate the level of importance of each domain. Colors for each domain are identical across panels.

For G2PCu (Figure 2C), which is influenced by the number of features contained in each network pair, 6 of the top 10 networks overlap with the LR-ENR and G2PCn results: SMN/SCN, SMN, VSN/SMN, CBN/VSN, CBN/SMN. G2Pcu results shed light on the importance of brain networks as a whole on the clustering without accounting for their size, which explains why the 4-ICN CBN network is not among the top 10 influential networks. Moreover, its comparison to G2PCn unveils the true nature of the importance of the features within each network (whether it is caused by individual feature importance or network size). For instance, CCN/SMN, which is formed by 17×9 ICN pairs, is ranked third in G2PCu and fifteenth in G2PCn. This implies that although CCN/SMN had an important effect on the clustering results, its underlying features were not as impactful as those of CBN (ranked first in the other methods). The combination of these methods helps identify bias towards larger networks and gives greater insight into the reasons behind the importance of each brain network.

### B. Identification of Differences in Cognitive Performance

We identified differences (i.e., p < 0.05) between the cognitive performance (i.e., FI2, NM1, PM1, PM2, and PM3) and demographics of each group (Table 1). G0 attempted fewer FI questions while obtaining a similar score, remembered more digits correctly, and took significantly less time to complete all pair matching rounds. Overall, G0 had better performance in visuo-spatial and reasoning tests. This widespread higher performance across cognitive tests could be related to a general intelligence factor discussed in previous studies [8].

**TABLE I.** Statistical Test Results

|  | Mann Whitney U-Tests |  |  |  |  |  |  | Chi-Squared Tests |  |
| --- | --- | --- | --- | --- | --- | --- | --- | --- | --- |
|  | NM1 | NM2 | FI1 | FI2 | PM1 | PM2 | PM3 | PrM1 | Sex |
| p-value | 0.25 | <0.001 | 0.24 | <0.001 | <0.001 | <0.001 | <0.05 | 0.93 | <0.001 |
| G0 > G1 | - | True | - | False | True | True | True | - | - |

While people in G0 attempted fewer fluid intelligence questions, there was no significant difference in their FI1 (correct number of questions), which implies a higher accuracy for answered questions. This could potentially be attributed to connectivity between the CBN and other networks [9]. NM1 records the total time spent taking the numeric memory test from the first to last round. In this test, the number of digits to be remembered increases with each correctly performed round. Although G0 remembered more digits than G1 (NM2), and thus completed more rounds, the time to complete the test was not significantly different, which implies that G0 memorized digits faster. Processing speed has been associated with the CBN [10].

Significant differences were found between the distributions of participant sex across clusters. G0 has a higher proportion of males than G1, which fits well with our clustering explainability results. Behavioral studies have shown that males and females have strong differences in intra-CBN connectivity and that males outperform females on spatial processing and motor and sensorimotor speed [11] that could be attributable to the SMN.

G0 performed faster in all PM rounds, which is potentially attributable to differences in intra-CBN FNC. The CBN has been shown to have an association with visual memory in patients with cerebellar ataxia [12]. Additionally, the CBN and thalamus in the SCN (i.e., CBN/SCN) play a role in motor coordination [13], which would be needed to perform the PM task. The SMN would also play a role in the coordination needed for the task, so a circuit between the CBN, CBN/SCN, SCN, SCN/SMN, SMN, and CBN/SMN, which were all important to the clustering, could play a significant role in the PM task.

### C. Limitations and Next Steps

While we identified relationships between the key networks identified with explainable clustering and existing literature related to cognitive performance, a useful next step would be to look for statistical relationships between the identified networks and the cognitive scores with differences between clusters. Also, due to the high data dimensionality (i.e., over 1,000 features), the clusters were not sensitive to the perturbation of individual features, and LR-ENR could not be counted upon to have used each feature in the same way as the original clustering. As such, we were somewhat restricted with the explainability approaches that we used and could only provide insight into the importance of each network connectivity pair as a result. It would be helpful to use more sensitive explainability approaches in the future as they are developed. Lastly, due to the large number of participants in the dataset, we used sFNC, rather than dFNC. Using an explainable clustering approach to identify dynamical patterns related to cognitive performance could be interesting.

## IV. Conclusion

In this study, we implement and compare several clustering explainability approaches. One approach, Normalized G2PC, is a novel variation on G2PC that provides better insight into the importance of brain network pairs. We apply the methods for insight into the brain network pairs that differentiate two subgroups of healthy individuals with different levels of cognitive function that we identify via clustering in a large sFNC dataset. We expect that our results will shed light on the role of brain network interactions in cognition and provide direction for integrating explainability methods into future rs-fMRI FNC clustering analyses.

